# Olfactory-to-visual facilitation in the infant brain declines gradually from 4 to 12 months

**DOI:** 10.1101/2023.09.08.556823

**Authors:** Diane Rekow, Jean-Yves Baudouin, Anna Kiseleva, Bruno Rossion, Karine Durand, Benoist Schaal, Arnaud Leleu

## Abstract

During infant development, intersensory facilitation declines gradually as unisensory perception improves. However, this developmental trade-off has been mainly investigated using audiovisual stimulations. Here, fifty 4- to 12-month-old infants were tested to determine whether the facilitating effect of their mother’s body odor on neural face categorization, as previously observed at 4 months, decreases with age. In a baseline odor context, results revealed a face-selective electroencephalographic (EEG) response that increases and changes qualitatively between 4 and 12 months, marking improved face categorization. At the same time, the benefit of adding maternal odor fades gradually with age, indicating an inverse relation with the strength of the sole visual response, and generalizing to olfactory-visual interactions previous evidence from the audiovisual domain.

## Introduction

From birth onward, human infants must navigate a complex multisensory environment and learn to form coherent percepts from a variety of sensory inputs. While the development of multisensory perception has long been debated (i.e., unisensory perception either preceding (Birch & Lefford, 1963; Piaget, 1952) or following (Gibson, 1969) multisensory perception), it is now generally admitted that infants can bind inputs across the senses at an early age, and that such intersensory integration improves and refines throughout development (Bahrick & Lickliter, 2012; Lewkowicz & Bremner, 2020; Murray et al., 2016 for reviews). Evidence accumulated so far indicates that the early emergence of multisensory perception is a function of principles that govern how inputs are integrated in the nervous system of numerous species.

A basic principle upon which infants rely to merge sensory inputs is *spatiotemporal coherence*. First described at the single-neuron level in nonhuman models (King & Palmer, 1985; Meredith et al., 1987), this principle ensues from the fact that the sensory features of an object do not occur arbitrarily, but are correlated in space and time. As a result, when auditory and visual stimuli are presented at same *vs*. different locations, human infants exhibit faster behavioral responses to stimuli coming from the same location already at 2 months of age (Neil et al., 2006). Likewise, when the temporal synchrony between a visual object and a sound is manipulated, neonates and infants aged up to 10 months evince distinct behavioral (Lewkowicz, 1996; Lewkowicz et al., 2010), autonomic (i.e., heart rate; Curtindale et al., 2019) and neural (Hyde et al., 2011; Werchan et al., 2018) responses to synchronous compared to asynchronous audiovisual stimulations. Strikingly, sensitivity to audiovisual synchrony is observed for both veridical (e.g., a ball bouncing up and down and a sound signaling the impact) and arbitrary (e.g., a human face and a tone) associations (Lewkowicz, 1996; Lewkowicz et al., 2010). This indicates a rudimentary ability to detect the synchronous onsets and offsets of concomitant inputs, forming a broadly tuned perceptual system that assembles simple events from their simultaneous occurrences in the physical environment (Murray et al., 2016).

Such early sensitivity to overlapping inputs has been proposed to support efficient perceptual learning, providing the scaffold on which more complex multisensory features can be integrated (Murray et al., 2016). This process would be particularly important during the initial learning of a specific domain, when intersensory correspondence between some features is still arbitrary for young infants (Lickliter & Bahrick, 2004, for review). In particular, the intersensory redundancy hypothesis (Bahrick & Lickliter, 2000) suggests that redundant information across several modalities (so-called amodal properties) grabs infants’ attention, renders inputs more salient and helps to bind features across the senses, leading to *intersensory facilitation* toward redundant information. For instance, during synchronous or collocated audiovisual stimulations, infants are able to discriminate tempos (at 3 months; Bahrick et al., 2002), rhythms (at 5 months; Bahrick & Lickliter, 2000), trajectories (at 4 months; Bremner et al., 2012), prosodies (at 4 months; Bahrick et al., 2019) or emotions (at 4 months; Flom & Bahrick, 2007), while they fail during unisensory, asynchronous or dislocated stimulations.

Interestingly, however, intersensory facilitation becomes less effective during development when unisensory perception improves. For instance, when exposed to audiovisual displays, young infants detect a change in the tempo and rhythm of a moving hammer at 3 and 5 months respectively, but they fail when the sole visual stimulus is presented (Bahrick et al., 2002; Bahrick & Lickliter, 2000). In contrast, older infants discriminate tempos and rhythms (at 5 and 8 months, respectively) from both unisensory and multisensory stimulations without any sign of intersensory facilitation (Bahrick & Lickliter, 2004). Remarkably, this developmental pattern can be reversed by task demand, as a more difficult tempo discrimination task leads 5-month-olds to perform like 3-month-olds, i.e., losing their discrimination ability with unisensory inputs and showing intersensory facilitation (Bahrick et al., 2010). Altogether, these findings are in line with another well-known principle of multisensory integration, *inverse effectiveness*, whereby the strength of the integration increases as unisensory responses decreases. First described in nonhuman models (Meredith & Stein, 1983), and later observed in human adults using behavioral (e.g., Regenbogen et al., 2016), neuroimaging (Stevenson & James, 2009) and electrophysiological (Stevenson et al., 2012) approaches, this principle can be transposed to multisensory development, as intersensory facilitation is particularly effective when unisensory perception is not fully developed.

So far, the principles subtending multisensory development were mainly investigated using auditory and visual stimuli, which are bound by a precise spatiotemporal synchrony. However, intersensory influences in early infancy can also occur for less space- and time-locked senses, such as olfaction (Sela & Sobel, 2010). For example, 3-month-old infants look longer at a smiling face associated with a pleasant rather than an unpleasant odor (Godard et al., 2016). At 4 months, exposure to the mother’s body odor increases looking duration at a face as opposed to a car (Durand et al., 2013), and at the mother’s face as opposed to a stranger’s face (Durand et al., 2020). At the same age, maternal odor facilitates the categorization of a variety of faces, as indexed by a larger face-selective electroencephalographic (EEG) response over the right occipito-temporal cortex (Leleu et al., 2020; Rekow et al., 2020). Similarly, when common objects configured as faces (i.e., facelike objects eliciting *face pareidolia* in adults) are rapidly presented among non-facelike objects belonging to the same categories, adding the mother’s body odor initiates a facelike-selective EEG response in 4-month-olds (Rekow et al., 2021). Later on, at 7 months, exposure to the mother’s odor also reduces the brain response to fearful faces (Jessen, 2020), and favors interbrain synchrony with, and visual attention to, an unfamiliar woman with whom the infant interacts (Endevelt-Shapira et al., 2021).

In most of these studies, odors were presented as contexts for long durations (i.e., several tens of seconds) and without clear spatial location. This indicates that despite a loose spatiotemporal relation between the two inputs, odors are prone to influence visual perception in infants. Does that mean that olfactory-to-visual facilitation does not rely on the same principles than other intersensory facilitations? Here, we address this question by investigating whether the maternal odor effect observed on neural face categorization at 4 months (Leleu et al., 2020; Rekow et al., 2020, 2021) follows the inverse effectiveness principle as applied to perceptual development, i.e., intersensory facilitation declining as unisensory perception develops. Indeed, the aforementioned studies used a rapid mode of visual stimulation together with a variety of naturalistic stimuli, making face categorization demanding for the 4-month-old brain. In adults, who effectively categorize genuine human faces from the sole visual input, there is no such improvement with a body odor, except for the less effective categorization of ambiguous facelike objects (Rekow, Baudouin, Durand, et al., 2022). Therefore, these findings suggest that the impact of a concurrent body odor progressively fades as the ability to categorize faces develops.

To tackle this issue, we used a cross-sectional design and a frequency-tagging EEG approach. We tested fifty infants, aged from 4 to 12 months, while they were exposed to fast streams of images presented at 6 Hz (6 images/s), with human faces inserted every 6^th^ image to tag a face-selective neural response at 1 Hz and harmonics (i.e., integer multiples) in their EEG spectra. During visual stimulation, infants were alternatively exposed to a maternal or a baseline odor context. In a first set of analyses, we examined the development of the face-selective response measured in the baseline odor context to determine whether face categorization becomes more efficient with age. In a second step, we analyzed the evolution of the maternal odor effect on face categorization as a function of age, hypothesizing an odor effect for the youngest infants that gradually disappears for the oldest infants. Finally, we conducted the same analyses on the general visual response to the fast train of images (6 Hz and harmonics) to assess whether the putative decline of the odor effect with age is selective to face categorization.

## Method

### Participants

Fifty-two healthy infants aged 4 to 12 months were recruited by mail from the local birth registry to participate in the study. Parents were informed about the objectives and methods of the study before they agreed to participate in signing a written informed consent. None indicated their infants having any sensory (e.g., visual, olfactory), neurologic, or psychiatric disorder. Procedure of testing was conducted according to the Declaration of Helsinki and approved by a national ethics committee ([blind for review]). One infant was excluded from the final sample due to less than two sequences per condition (see below) and another infant because of too noisy EEG data. The final sample was thus composed of 50 infants (26 females) whose age covered a large range (121–374 days, mean age ± SD: 242 ± 78 days, Figure 1A). Sample size was estimated a priori by considering a moderate relation between the age of the infants and the maternal odor effect (*R*² = 0.20), a significance level *α* = .05 (two-tailed), and a power 1-*β* = .90, leading to a sample size *N* = 50.

**Figure 1.**
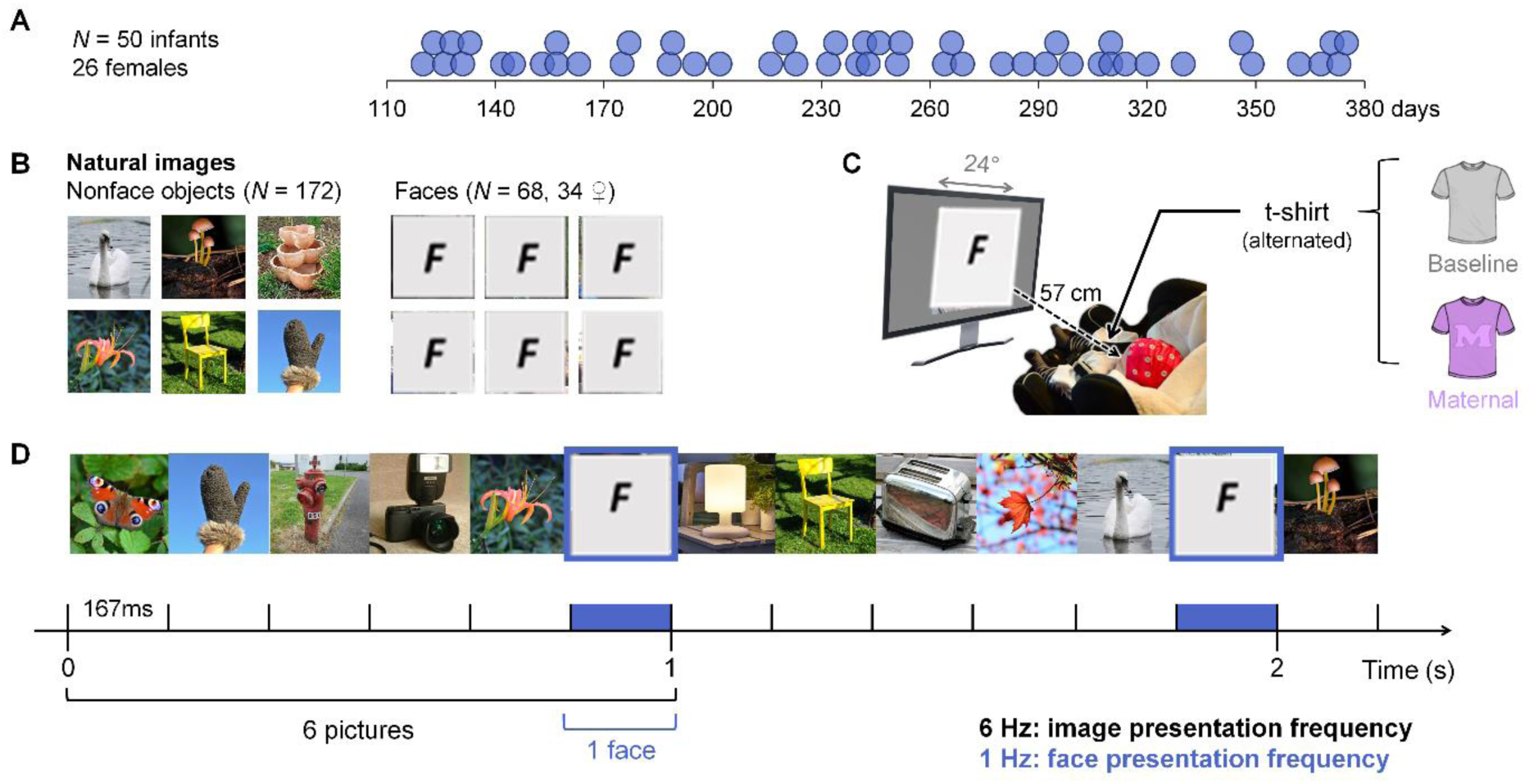
Participants, stimuli and procedure. **A.** Age distribution of the 50 included infants. Circles represent individual participants as a function of their age (in days) at the time of testing. **B.** Examples of the natural images of control objects and faces (here replaced by “F” placeholders for copyright issues) used as stimuli. **C.** Infants were equipped with an electrode cap and seated at 57 cm in front of the stimulation screen. Before a series of 2 sequences of visual stimulation (36 s each), a folded white t-shirt imbued with the odor (either maternal, depicted in violet, or baseline, depicted in grey) was placed on their chest. T-shirts were alternated every 2 sequences. **D.** Extract of 2.167-s of the 34.5-s-long clip of visual stimulation. Each stimulus lasted 167 ms and was directly followed by another stimulus, resulting in a fast 6-Hz rate of image presentation (6 images/s). A face (F) was presented every 6^th^ stimulus, thus corresponding to a 1-Hz face presentation frequency (1 face/s).

### Visual stimuli

Natural colored images of various nonface objects (animals, plants, man-made objects; N = 172) and human faces (N = 68, 34 females) were used (Figure 1B). Images were cropped to a square and resized to 400 × 400 pixels. In the cropped images, items appeared at variable locations and the original background was preserved, so that physical cues were highly variable across stimuli and barely informed about the visual category. Stimuli were presented in the center of a 24-inch LED screen (refresh rate: 60 Hz, resolution: 1920 × 1080 pixels, background: 128/255 in grayscale) at a viewing distance of 57 cm, subtending 24 × 24° of visual angle.

### Odor stimuli

Following previous studies, two odor stimuli were used: the mother’s body odor and a baseline odor (Durand et al., 2013; Leleu et al., 2020; Rekow et al., 2020, 2021). Both odors were delivered using white cotton t-shirts. T-shirts were first laundered using a scentless hypoallergenic powder detergent (Persavon, France). One t-shirt was sent to the mother of each tested infant in a zip-locked hermetic plastic bag. Instructions were enclosed for the collection of her body odor, specifying to wear the t-shirt on bare skin during the three consecutive nights preceding the experiment and to refrain from using odorous soap or perfume before wearing it. During the collection period, mothers had to store the t-shirt in the hermetic bag at room temperature (away from any heating device). The odor of an unworn t-shirt (stored in a similar plastic bag in our premises) was used as the baseline odor.

### Procedure

Procedure was identical to that of the aforementioned infant frequency-tagging EEG studies (Leleu et al., 2020; Rekow et al., 2020, 2021). Testing took place in a light- and sound-attenuated room equipped with an air-renewing system. To additionally reduce olfactory noise, the room was aired between testing sessions and experimenters did not use odorous products prior to, or during, the session. Each infant was equipped with an electrode cap and seated in a baby car seat in front of the screen behind occluding blinds to minimize visual distractions. Parents were asked to stay at a minimal 2.5-m distance away from their infant and to refrain from interacting with them except in case of manifest distress. A webcam allowed to monitor the infant and to launch the visual stimulation when the infant looked at the screen. The t-shirts were manipulated by the experimenters using dedicated disposable nitrile gloves (Schield Scientific, The Netherlands). To deliver the odors, they were folded backward (sleeves folded over the front) to expose the infant to the breast and axillary areas, and disposed on their upper chest while being maintained by the seat belts (Figure 1C). One t-shirt corresponding to one odor condition was placed just before the beginning of a sequence of visual stimulation (see below), and the two odor conditions were alternated every two sequences. Odor presentation order was counterbalanced across infants, and a minimum interval of 1 min was introduced at each change, taking the form of a break of visual stimulation for the infant.

Sequences of periodic visual stimulation were presented at a fast rate of 6 Hz (6 images/s) without inter-stimulus interval, such that each stimulus lasted 167 ms (i.e., 1 s/6) on the screen. Face images were inserted among nonface stimuli as every 6^th^ stimulus, i.e., at a rate of 6 Hz/6^th^ = 1 Hz (i.e., 1-s interval between each face, Figure 1D). As a result, we tagged two distinct neural responses at different frequencies in the EEG amplitude spectrum: a general visual response at 6 Hz and harmonics (i.e., integer multiples) and a face-selective response at 1 Hz and harmonics. The general visual response comprises the neural activity elicited by the rapid change of stimuli 6 times per second, and is mainly sensitive to the variations of low-level physical cues (e.g., local cues, contrast). In contrast, the face-selective response captures the neural activity selectively elicited by the face images, i.e., a direct differential response to faces against many living and non-living categories, insensitive to low-level cues (already captured by the general visual response) and generalized across variable individual faces.

Each sequence of visual stimulation lasted 34.5 s and was composed of a pre-stimulation interval (0.5 s), a fade-in of ramping-up contrast (0 to 100%, 1.833 s), the full-contrast stimulation segment (31.167 s), a fade-out of decreasing contrast (100% to 0, 0.833 s) and a post-stimulation interval (0.167 s). The 68 face images were randomly split into two sets of 34 faces (17 females) and counterbalanced between sequences, therefore all individual faces were presented equally across the two consecutive sequences of one odor condition, and contrasted to the same set of 172 nonface stimuli. If needed, short sounds were used to reorient infant’s attention to the screen (sporadic and non-periodic, thus not contaminating the frequency-tagged EEG responses with auditory-evoked potentials). The experiment stopped when the infant manifested disinterest or fatigue, or at parental demand. Infants were included in the final sample if they achieved at least 2 valid sequences per odor condition (i.e., no premature abortion and presence of a general visual response, see next section).

### EEG recording and preprocessing

EEG was recorded from a 32-Ag/AgCl-electrode cap (Waveguard, ANT Neuro, The Netherlands) according to the 10–10 classification system (acquisition reference: AFz, electrode initial impedance < 15 kΩ, sampling rate: 1024 Hz). EEG analyses were run on Letswave 6 (http://www.letswave.org/) carried out using Matlab 2017 (MathWorks, USA). Left and right mastoid electrodes (M1 and M2) were too noisy and thus removed from montage before preprocessing. In a first step, individual EEG data were preprocessed and cleaned of artifacts, blind of the condition (see Supplementary Method for details). After preprocessing, the final number of epochs ranged between 2 to 8 per infant, with an overall rejection of 18 epochs out of 398 (i.e., 4,5%). The remaining average number of epochs was (mean ± SD) 3.64 ± 1.52 and 3.74 ± 1.65 for the baseline and maternal odor conditions, respectively. After this cleaning step, remaining epochs were sorted according to the odor condition and averaged together in the time domain to obtain a single 32-s-long epoch per condition for each infant. These data are made available in the public repository associated with this manuscript (https://osf.io/twyp5/?view_only=ccada0e1575046499fae51301f08afdc).

### Frequency-domain analysis

A fast Fourier transform was applied to the two epochs (one per condition) for each infant and raw amplitude spectra were extracted for all electrodes with a frequency resolution of 1/32 = 0.03125 Hz. Given the steep power-law function of the EEG spectrum, a baseline correction was first applied to remove background noise and lead to a notional amplitude of zero in the absence of frequency-tagged responses. At each frequency bin, noise was defined as the mean of 6 neighboring bins and subtracted out. These bins were selected among the 10 surrounding bins (5 on each side: ± 0.15625 Hz) after the exclusion of the 2 immediately adjacent (one on each side, in case of spectral leakage) and the 2 extreme (minimum and maximum) bins (to avoid including signal and potential outliers in noise estimation). Next, we estimated the range of significant harmonics separately for the general visual response (6 Hz and integer multiples) and the face-selective response (1 Hz and integer multiples) using *Z*-scores calculated on the average of all electrodes, infants and odor conditions. Z-scores were computed as the difference between the amplitude at the frequency of interest and the mean amplitude of 20 neighboring bins, divided by their standard deviation. These bins were selected among 22 surrounding bins beyond those used for baseline correction (11 on each side: from ± 0.1875 Hz to ± 0.5 Hz) after the exclusion of the two extreme bins. Harmonics were considered significant when their *Z*-score was > 1.64 (*p* < .05, one-tailed, signal > noise). The range of significant harmonics was defined until *Z*-scores were no longer significant. For the general visual response, 6 consecutive harmonics were significant (i.e., until 36 Hz, all *Z*s > 5.06, Table S1), and significance reached the 7^th^ harmonic for the face-selective response (i.e., 7 Hz, all *Z*s > 2.15, Table S1), excluding the 6^th^ harmonic which corresponds to the general visual response (i.e., 6 Hz).

Electrodes of interest were then identified on the responses summed across significant harmonics and still averaged across infants and odor conditions. For each brain response, the *Z*-scores of all individual electrodes were calculated and considered significant when *Z* > 2.93 (*p* < .05, one-tailed; signal > noise, Bonferroni-corrected for 30 electrodes). For the general visual response, every individual electrode showed a significant response (all *Z*s > 17.6, Table S2). Therefore, we kept the 4 best electrodes that were all located over the middle occipital cortex (Oz: *Z* = 168, O2: *Z* = 133, POz: *Z* = 122 and O1: *Z* = 102). For the face-selective response, 11 electrodes showed a significant response (all *Z*s > 3.20, Table S2). We considered the 4 best electrodes excluding the midline, which corresponded to 2 left and 2 right homologous occipito-temporal electrodes (P8: *Z* = 19.5, P7: *Z* = 13.1, O1: *Z* = 5.31 and O2: *Z* = 4.38). For both responses, electrodes of interest matched those reported in previous frequency-tagging EEG studies investigating face categorization in infants (de Heering & Rossion, 2015; Leleu et al., 2020; Rekow et al., 2021).

Subsequent analyses were completed for both the face-selective response and the general visual response, but in separate pipelines. In a first step, the development of each neural response as a function of age was characterized in the baseline odor context. A repeated-measures ANCOVA was run on individual amplitudes with *Age* as a continuous factor, and *Electrode* (either P7, P8, O1, O2 for the face-selective response or POz, Oz, O1, O2 for the general visual response) and *Harmonic* (1^st^, 2^nd^, 3^rd^, 4^th^, 5^th^ and either 6^th^ for the general visual response or 7^th^ for the face-selective response) as within-subject categorical factors. *F* values and partial eta squared (*η*_p_²) are reported, and significance threshold was fixed at *p* < .05. Sphericity was assessed using Mauchly’s test and whenever it was violated, the Greenhouse-Geisser correction was applied (corresponding effects are reported with adjusted degrees of freedom and epsilon coefficient *ε*). Given our aim to characterize the development of the neural responses, we focused on the main effect of *Age* and its interactions with the other factors. Contrasts were used to decompose significant interactions. Then, another repeated-measures ANCOVA with the *Age* and *Electrode* factors was conducted on individual amplitudes summed across harmonics with (a) significant effect(s) involving *Age*. Finally, to assess the evolution of the number of significant harmonics as a function of age, *Z*-scores, as defined above, were calculated for each infant, electrode and harmonic, considering the 4 electrodes and the 6 harmonics defined at group level for each response. The maximum number of significant harmonics (i.e., *Z* > 1.64, *p* < .05, one-tailed, signal > noise) for at least one electrode was extracted for each infant and submitted to a linear regression with *Age* as a continuous factor. *R*² and *F* values are reported, and significance threshold was fixed at *p* < .05. For all these analyses, the effect of *Age* is illustrated with the predicted outcomes from the regression line equation at X = 120 days (4 months) and X = 360 days (12 months), and with the data averaged for the 25 youngest infants (4-8 months) and the 25 oldest infants (8-12 months). Means and standard errors of the means (SEM) are thus reported for these two subgroups but were not submitted to significance testing.

In a second step, we analyzed the difference between the two odor conditions as a function of age. As for the previous analysis, a repeated-measures ANCOVA was first computed on individual amplitudes using the *Age*, *Electrode* and *Harmonic* factors, with the addition of *Odor* (maternal, baseline) as a within-subject categorical factor. Again, *F* values and *η*_p_² are reported, the Greenhouse-Geisser correction was applied whenever necessary, and significance threshold was fixed at *p* < .05. Here, given our aim to determine whether maternal odor exerts an influence on the visual responses and, if so, whether this effect declines as age increases, we focused on effects involving the *Odor* factor and its interaction with the *Age* factor. Significant interactions were decomposed using contrasts. As for the first step of analyses, another repeated-measures ANCOVA with the *Age*, *Electrode* and *Odor* factors was conducted on individual amplitudes summed across relevant harmonics. For illustration purposes, predicted outcomes of the *Odor* × *Age* interaction are reported together with data averaged for the youngest and oldest infants (see above). Finally, we computed a lateralization index by considering raw amplitudes summed across harmonics at electrodes O1 and O2 (since O2 is the only electrode showing a significant decrease of the odor effect with age, see Results). Amplitude at O1 was subtracted from amplitude at O2 and then divided by the sum of the two electrodes to reflect the advantage for one hemisphere expressed in %, with positive and negative values corresponding to right- and left-lateralized responses, respectively. We calculated the index for each infant and each odor context and submitted individual odor effects (maternal minus baseline odor) to a linear regression with *Age* as a continuous factor.

## Results

### Face-selective neural activity progressively increases and refines with age

To delineate the development of the ability to rapidly categorize human faces within a fast train of natural images between 4 and 12 months of age, a first set of analyses was conducted in the baseline odor context. Visual inspection of the EEG spectrum revealed that the periodic appearance of face stimuli at 1 Hz within the rapid visual stimulation elicits a clear response at the same frequency and harmonics over the occipito-temporal cortex for both the youngest (4-8 months) and the oldest (8-12 months) infants. Descriptively, this face-selective response is larger for the oldest infants for the 1^st^, 2^nd^, 4^th^ and 5^th^ harmonics, while lower for the 3^rd^ and 7^th^ harmonics. Summed across these 6 harmonics, the overall response is larger (+128%) for the oldest (2.60 ± 0.47 µV) than the youngest (1.14 ± 0.31 µV) infants (Figure 2A).

**Figure 2.**
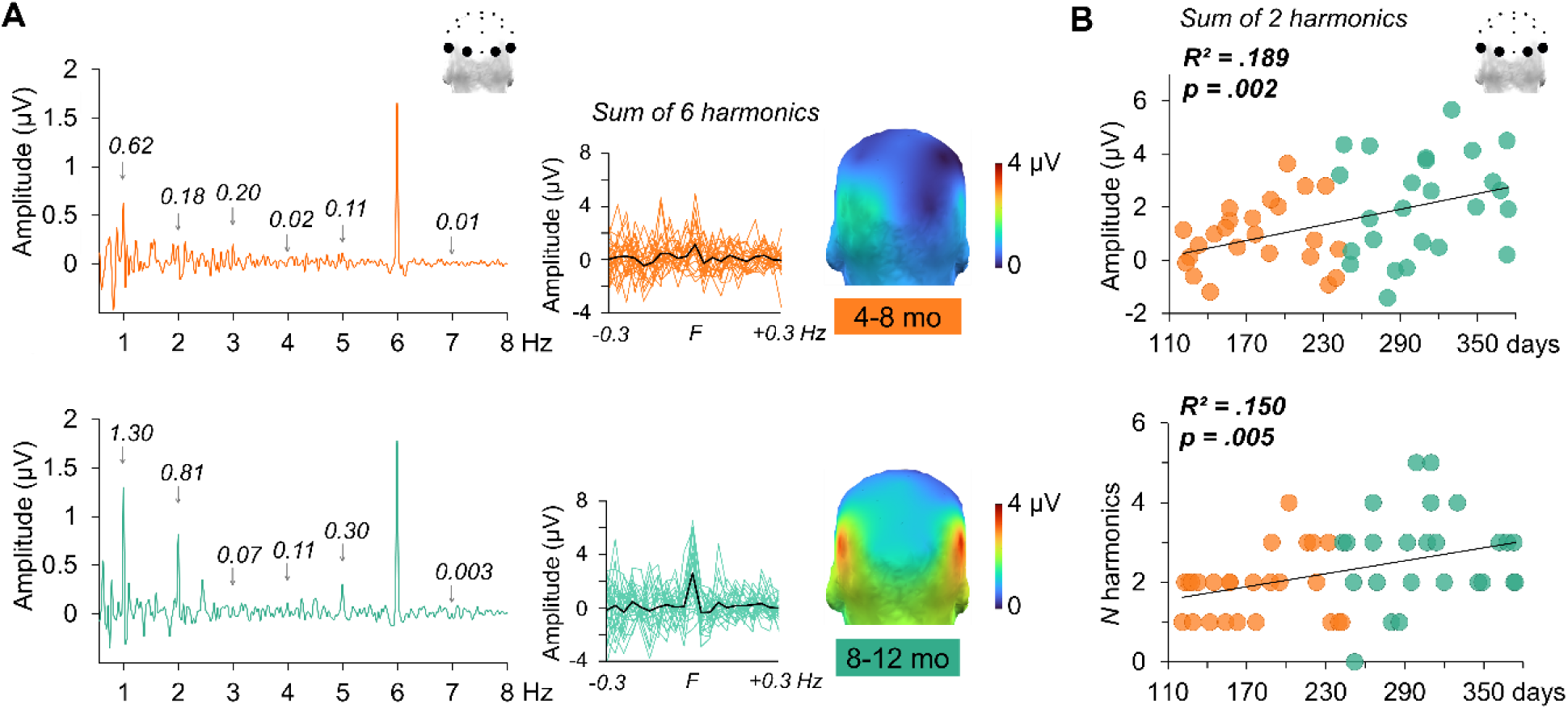
The development of the face-selective response with age. **A.** Amplitude spectra recorded in the baseline odor context for the youngest (orange: 4-8 months, top) and the oldest (green: 8-12 months, bottom) infants averaged for the 4 occipito-temporal electrodes (P7, O1, O2, P8). The face-selective response is captured at 1 Hz and harmonics (until 7 Hz) excluding the 6^th^ harmonic (6 Hz) which corresponds to the general visual response. The smaller spectra represent the summed amplitude of the response, with the average group-level activity in black and individual spectra in color. Head maps show the topography (posterior view) of the summed response. **B.** Amplitude of the face-selective response summed across the 2 first harmonics (top) and maximum number of significant harmonics (bottom) as a function of age. Each circle represents individual infant data depending on their subgroup (orange: 4-8 months, green: 8-12 months).

The analysis of individual amplitudes extracted for each harmonic at occipito-temporal electrodes (P7, P8, O1, O2) revealed a significant main effect of *Age* (*F* (1, 48) = 8.41, *p* = .006, *η_p_*² = .149), qualified by a significant *Harmonic* × *Age* interaction (*F* (1.9, 90.3) = 5.54, *p* = .006, *η_p_*² = .104, *ε* = .38). A significant effect of *Age* was found for the 1^st^ (*F* (1, 48) = 6.03, *p* = .018, *η_p_*² = .112) and 2^nd^ (*F* (1, 48) = 12.01, *p* = .001, *η_p_*² = .200) harmonics, the face-selective response increasing as a function of age for both of them (Figure S1). No effect of *Age* was found for the other harmonics (all *F*s < 2.18, *p*s > .14). The analysis on the sum of the two first harmonics also yielded a significant main effect of *Age* (*F* (1, 48) = 11.17, *p* = .002, *η_p_*² = .189). The predicted face-selective response combined across these harmonics is 10 times larger at 12 months (i.e., 360 days: 2.61 µV) than 4 months (i.e., 120 days: 0.26 µV) (Figure 2B left), and its mean amplitude measured for the youngest infants (4-8 months) is 0.80 ± 0.28 µV, as opposed to 2.11 ± 0.37 µV for the oldest infants (8-12 months). In sum, there is a strong increase of the face-selective response between 4 and 12 months that is mainly driven by the two first harmonics. The outcomes of these ANCOVAs are reported in Table S3.

We also determined whether the number of significant harmonics evolves between 4 and 12 months. We extracted the maximal range of significant harmonics for each infant (Figure 2B right), and found an effect of *Age* (*R*² = .150, *F* (1, 48) = 8.48, *p* = .005) showing that the number of harmonics increases significantly as age increases (predicted number: from 1.62 harmonics at 120 days to 2.92 harmonics at 360 days). On average, the youngest infants (4-8 months) have 1.88 ± 0.17 significant harmonics compared to 2.68 ± 0.18 for the oldest infants (8-12 months). Overall, all infants but one (i.e., 98% of infants) present at least 1 significant harmonic. Hence, the face-selective response does not only increase with age but also complexifies, being distributed on more harmonics in the oldest infants.

### The influence of maternal odor on the face-selective response gradually declines with age

The second objective was to estimate whether the influence exerted by maternal odor on the face-selective response evolves with age. In line with a previous study conducted at 4 months (Leleu et al., 2020), visual inspection of topographical head maps suggested a larger response over the right occipito-temporal cortex in the presence of the mother’s body odor compared to the baseline odor, especially for the youngest 4-8-month-old infants (Figure 3A). Indeed, their face-selective response increases with maternal odor at both electrodes O2 (from -0.19 ± 0.34 µV in the baseline odor context to 1.12 ± 0.37 µV in the maternal odor context) and P8 (from 1.06 ± 0.45 µV to 1.76 ± 0.40 µV). In contrast, for the oldest infants (8-12 months), the response increases only at P8 (from 3.42 ± 0.74 µV to 4.20 ± 0.94 µV) and decreases at electrode O2 (from 1.06 ± 0.39 µV to 0.13 ± 0.38 µV). In addition, for both age groups, the face-selective response decreases with maternal odor at the two left-hemispheric electrodes, although to a lesser extent than the right-hemispheric increase (mean across O1 and P7, respectively from 1.16 ± 0.30 µV in the baseline odor context to 0.78 ± 0.33 µV in the maternal odor context at 4-8 months and from 1.97 ± 0.41 µV to 1.58 ± 0.43 µV at 8-12 months).

**Figure 3.**
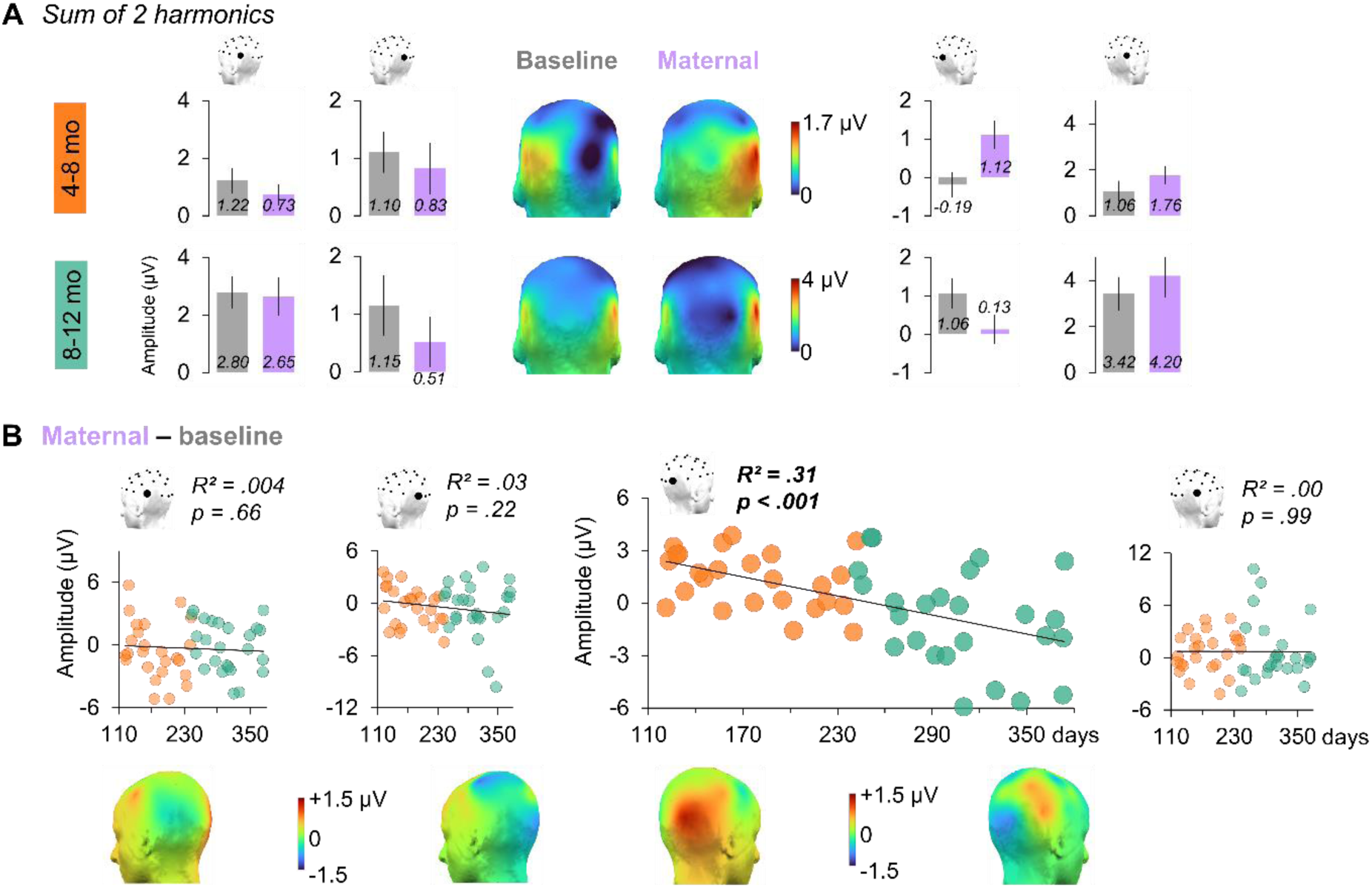
Maternal odor effect on the face-selective response. **A.** Amplitude of the face-selective response summed across two harmonics at each occipito-temporal electrode (from left to right: P7, O1, O2, P8) for the youngest (4-8 months, top) and the oldest (8-12 months, bottom) infants in the baseline (grey) and maternal (violet) odor contexts. Head maps show the topography (posterior view) of the response in each odor context and for each subgroup. **B**. Maternal odor effect (amplitude of the face-selective response in the maternal minus the baseline odor context) at each occipito-temporal electrode as a function of age. Each circle represents individual infant data depending on their subgroup (orange: 4-8 months, green: 8-12 months). Below are head maps showing the topography (lateral views) of the effect for the youngest (4-8 months) and the oldest (8-12 months) infants.

We analyzed individual amplitudes extracted for each harmonic and each odor context and found a marginal *Harmonic* × *Odor* × *Age* interaction (*F* (2.3, 110.8) = 2.65, *p* = .067, *η_p_*² = .052, *ε* = .46) qualified by a significant *Harmonic* × *Odor* × *Electrode* × *Age* interaction (*F* (5.0, 241.7) = 4.28, *p* < .001, *η_p_*² = .082, *ε* = .34). Decomposition of this interaction revealed that the *Odor* effect and its modulation by *Age* are limited to the two first harmonics, since no other harmonics evinced a significant effect involving these factors (all *F*s < 2.50, *p*s > .081). For the 1^st^ harmonic, we found significant *Odor* × *Electrode* and *Odor* × *Electrode* × *Age* interactions (both *F*s (3, 144) > 4.31, *p*s < .006, *η_p_*² > .083). For the 2^nd^ harmonic, there was a significant *Odor* × *Age* interaction (*F* (1, 48) = 6.12, *p* = .017, *η_p_*² = .113).

When the face-selective response is summed across these two harmonics, the analysis yielded a significant main effect of *Odor* (*F* (1, 48) = 5.15, *p* = .028, *η_p_*² = .097) and several interactions including the *Odor* factor, especially the *Odor* × *Electrode* × *Age* interaction (*F* (3, 144) = 3.35, *p* = .021, *η_p_*² = .065). Contrasts indicated that the *Odor* × *Age* interaction is actually limited to the right occipital electrode O2 (*F* (1, 48) = 21.83, *p* < .001, *η_p_*² = .313), which shows a predicted increase of amplitude of +2.40 µV in the maternal *vs*. the baseline odor context at 4 months (120 days), compared to a reduction of -1.95 µV at 12 months (360 days) (Figure 3B). The youngest infants (4-8 months) have a mean odor effect of +1.31 ± 0.32 µV as opposed to -0.93 ± 0.56 µV for the oldest infants (8-12 months). The three other electrodes displayed neither the main effect of *Odor* nor its modulation by *Age* (all *F*s < 1.58, *p*s > .21), despite a positive odor effect on average for P8 (+0.74 ± 0.42 µV) and negative odor effects for P7 and O1 (-0.32 ± 0.36 µV and -0.46 ± 0.38 µV, respectively). The outcomes of the ANCOVAs are reported in Table S4.

Finally, to further confirm the lateralization of the maternal odor effect as a function of age, we computed a lateralization index between O1 and O2 (with positive and negative values corresponding to right and left hemisphere advantages, respectively) and calculated individual odor effects (maternal minus baseline). We found a significant effect of *Age* (*R*² = .103, *F* (1, 48) = 5.54, *p* = .023) confirming the increase of the face-selective response over the right hemisphere with the mother’s body odor at 4 months (predicted index at 120 days: +24.9%) that progressively declines (predicted index at 360 days: -7.5%). The mean odor effect on the index is +17.1 ± 5.4% for the youngest infants (4-8 months) as opposed to -0.3 ± 7.2% for the oldest infants (8-12 months). Hence, in essence, the presence of the mother’s body odor strongly increases the face-selective response recorded over the right occipital cortex at 4 months, this odor effect gradually declining with age. At 12 months, the odor effect reverses, the face-selective response decreasing over the right occipital region to become restricted to more anterior occipito-temporal locations.

### No change of the general visual response with age and in the presence of maternal odor

The same analysis pipeline as for the face-selective response was applied to the general visual response to the rapid stream of natural images (6 Hz and harmonics). Inspection of the EEG spectrum recorded in the baseline odor context indicated a clear response at 6 Hz and harmonics over the middle occipital cortex at all age (Figure S2A). This response is descriptively larger for the youngest (4-8 months) than the oldest (8-12 months) infants for the 2^nd^, 3^rd^ and 4^th^ harmonics and lower for the 1^st^, 5^th^ and 6^th^ harmonics, leading to an overall response (summed across the 6 harmonics) of 4.36 ± 0.57 µV for the youngest infants and 4.15 ± 0.65 µV for the oldest infants (Figure 5A). The general visual response is not different between the two odor contexts for the youngest infants (mean amplitude in the maternal odor context: 4.36 ± 0.61 µV) while slightly larger with the mother’s body odor for the oldest infants (4.67 ± 0.71 µV).

**Figure 4.**
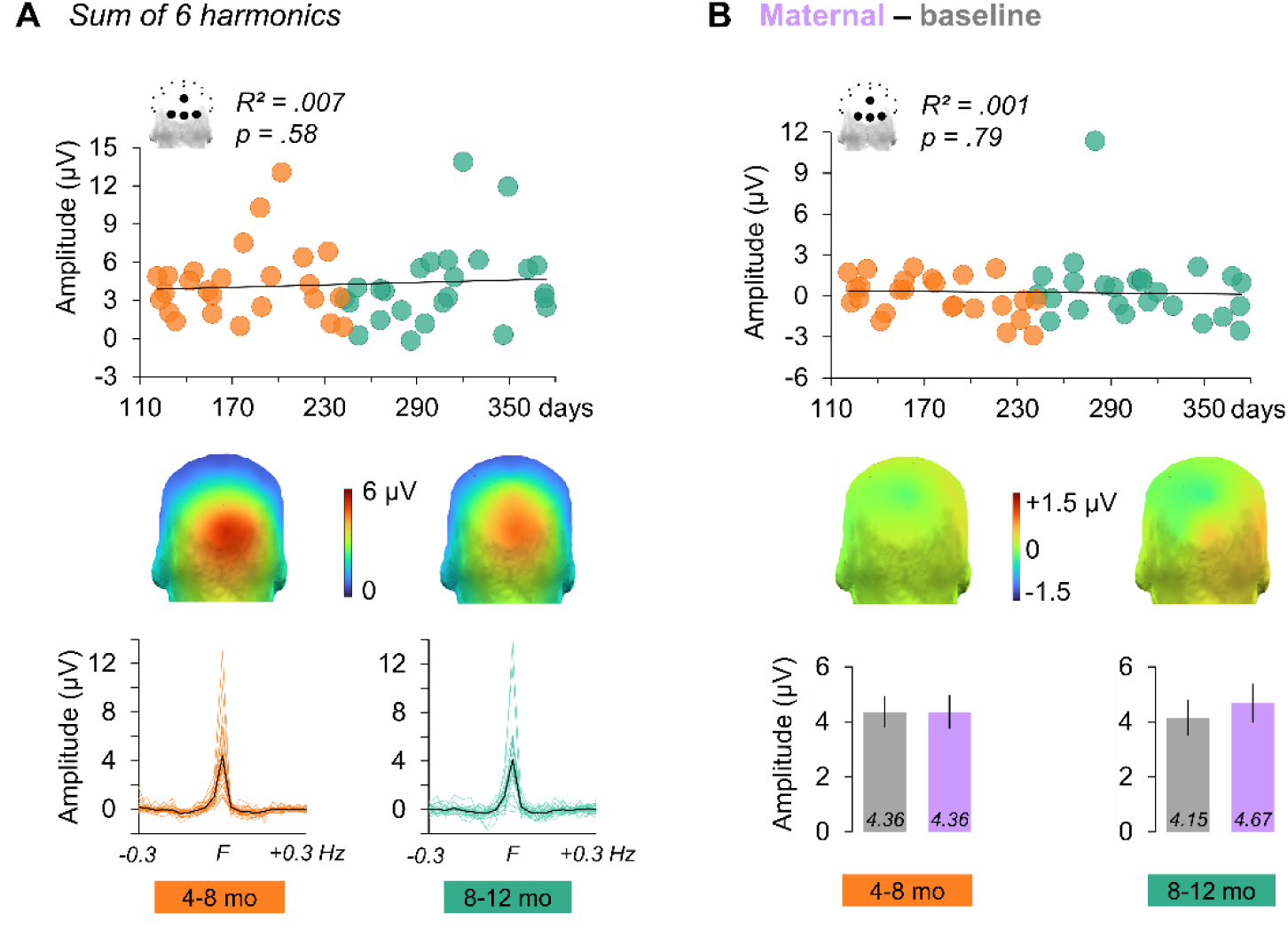
No effect of age and maternal odor on the general visual response. Amplitude of the general visual response averaged for the 4 medial occipital electrodes (Oz, POz, O1, O2) and summed across six harmonics (from 6 to 36 Hz) as a function of age in the baseline odor context (**A**) and for the difference between the maternal (violet) and the baseline (grey) odor contexts (maternal odor effect, **B**). Each circle represents individual infant data depending on their subgroup (orange: 4-8 months, green: 8-12 months). Head maps show the topography (posterior view) of the response. The small spectra in A represent the mean response of each subgroup in black and individual spectra in color. Bar graphs in B depict the mean amplitude of the response for each subgroup and each odor context.

The analysis of individual amplitudes extracted for each harmonic at medial occipital electrodes (O1, O2, Oz, POz) in the baseline odor context did not reveal a significant effect of *Age* (*F* (1, 48) = 0.32, *p* = .58, *η_p_*² = .007, Figure 5A) or any interaction involving this factor (all *F*s < 1.90, *p*s > .16). Similarly, the maximum number of significant harmonics remains stable as a function of age (mean number across infants: 5.0 ± 0.16 significant harmonics), as revealed by a non-significant linear regression (*R*² = .009, *F* (1, 48) = 0.45, *p* = .51, Figure S2B). Finally, the analysis including the *Odor* factor did not reveal a main effect or interaction with this factor (all *F*s < 0.46, *p*s > .60), for a mean odor effect (maternal minus baseline) across infants of +0.26 ± 0.30 µV for the summed response across harmonics (Figure 5B). Since one infant has an odor effect of +11.3 µV whereas the remaining 49 infants have an effect comprised between -2.96 µV and 2.46 µV, we reran the analyses without this outlier. The updated analysis did not change the conclusions (all *F*s < 0.98, *p*s > .32), for a mean odor effect across infants of +0.04 ± 0.20 µV. The outcomes of these ANCOVAs are reported in Table S5.

## Discussion

By using frequency-tagging EEG to measure rapid face categorization in fifty infants aged from 4 to 12 months, we hereby identify that a face-selective neural response recorded over the occipito-temporal cortex develops as a function of age, both quantitatively (larger amplitude) and qualitatively (distributed on more harmonics in the EEG spectrum). Most importantly, by exposing the infants to their mother’s body odor or a baseline odor, we also replicate previous evidence of a larger face-selective response over the right hemisphere in the presence of the mother’s odor for the youngest infants (Leleu et al., 2020; Rekow et al., 2021), and further demonstrate that this maternal odor effect gradually declines as face categorization develops (with age). Critically, the general response to the rapid stream of visual stimulation recorded over the middle occipital cortex is immune to the presence of maternal odor, excluding a mere influence on visual attention or general arousal. Overall, we provide conspicuous evidence that during early perceptual development, intersensory facilitation between olfaction and vision decreases as visual perception develops, extending previous findings with audiovisual stimulations (Bahrick et al., 2004) and generalizing them to olfactory-visual interactions.

### The development of rapid face categorization between 4 and 12 months

A first major achievement of the present study is to delineate the development of rapid face categorization in the infant brain between 4 and 12 months. Face perception in general follows a well-documented protracted development in infancy (Pascalis et al., 2020), several abilities improving during the first year of life, such as the discrimination of facial identities (Sugden & Marquis, 2017, for a meta-analysis) or expressions (e.g., Poncet et al., 2022 for recent evidence with frequency-tagging EEG). At birth, infants already orient toward simple face-like stimuli (Johnson et al., 1991), an inborn ability that may be driven by an early preference for basic configurations of visual features (Simion et al., 2007 for a review). Then, between 3 and 4 months, infants exhibit preferential looking for a face over one other object (e.g., a car; de Heering et al., 2016; Durand et al., 2013). Similarly, several EEG studies have measured the event-related potentials elicited by face *vs.* nonface stimuli and identified distinct neural activity for faces from 3-4 months (see de Haan et al., 2003 for review; see also Conte et al., 2020, for recent evidence). However, when testing infants in more complex visual settings, such as a face presented among several nonface objects (Di Giorgio et al., 2012; Kwon et al., 2016) or in more naturalistic stimuli (Frank et al., 2014; Kelly et al., 2019), rapid face detection as measured by eye-tracking mainly improves from 6 months of age (Leppänen, 2016 for review), highlighting that younger infants difficultly track faces in rich environments. At the neural level, previous infant studies have also used frequency-tagging EEG to measure the rapid (i.e., at a glance) categorization of faces contrasted to many other living and nonliving nonface categories within a set of highly variable naturalistic stimuli (de Heering & Rossion, 2015; Leleu et al., 2020). They have identified an occipito-temporal face-selective response already at 4-6 months, which is relatively small (about 1 µV), focal and captured in a single harmonic. Similar studies in children and adults have shown that the face-selective response becomes larger and distributed on several harmonics at 5 (Lochy et al., 2019) and 10 years of age (Vettori et al., 2019), before being observed on even more harmonics with a large occipito-temporal topography in adults, for an overall amplitude of about 4 µV (e.g., Jacques et al., 2016; Rossion et al., 2015). Thus, extending these studies, here we reveal that the frequency-tagged face-selective response progressively increases between 4 and 12 months, with a predicted amplitude of 0.26 µV at 4 months to 2.61 µV at 12 months. Moreover, though here the bulk of the response concentrates on the two first harmonics, which both show a strong effect of age, the number of significant harmonics also increases between 4 and 12 months (predicted number of 1.62 at 4 months to 2.92 at 12 months), indicating that the brain response complexifies (multicomponent waveform in the time domain, see Retter et al., 2021), despite a much lower number of harmonics than in adults (e.g., Jacques et al., 2016; Rossion et al., 2015). Hence, altogether, these findings demonstrate that the ability to rapidly categorize a variety of faces within naturalistic stimuli follows a gradual development during the first year. Future studies should pursue these efforts to delineate the development of face-selective neural activity beyond 1 year of age.

Remarkably, the qualitative and quantitative changes of the face-selective response between 4 and 12 months may reflect improved categorization abilities for both faces and nonface items. In frequency-tagging paradigms of rapid categorization (e.g., Barbero et al., 2021; Jacques et al., 2016; Rekow, Baudouin, Brochard, et al., 2022; Rekow, Baudouin, Durand, et al., 2022), the category-selective response is a direct differential response reflecting the discrimination of the target category from the other categories displayed in the sequence (i.e., distinct neural activity), and the generalization of this discrimination to different exemplars of the target category (i.e., reliable distinct neural activity during the whole sequence). Hence, the observed larger amplitude of the response as age increases, suggests that infants become better at discriminating faces from other objects and at generalizing this response across the variable natural images of faces, which depend on both the accurate recognition of face stimuli (high rate of hits among the 34 faces within a sequence) and the absence of face-selective activity for nonface stimuli (low rate of false alarms among the 172 nonface objects). In other words, improved face categorization does not only imply that faces become more familiar to infants and have a higher chance to be perceived as faces, but also that nonface objects become more familiar to infants and have a lower chance to be *misperceived* as faces. Finally, it is worth noting that this developmental pattern is specific to the face-selective response, as the general visual response does not change with age, in line with previous studies at various ages (de Heering & Rossion, 2015; Leleu et al., 2020; Lochy et al., 2019; Vettori et al., 2019). Given that the general response reflects basic visual function in response to the fast train of images (i.e., detection of changes between 2 successive stimuli), this suggests a selective perceptual improvement based on infants’ experience and irrespective of the mere maturation of the visual system, likely driven by the sheer ubiquity of faces in the infants’ environment (Jayaraman et al., 2015).

### The mother’s body odor fosters face categorization in the youngest infants

A second major achievement of the study is to replicate the maternal odor effect on face-selective neural activity previously found at 4 months for both genuine human faces (Leleu et al., 2020) and naturalistic face-like stimuli (Rekow et al., 2021). In line with these studies, the effect is observed over the right occipital cortex, dominant for face categorization (Rossion & Lochy, 2021 for review). Other studies have also found that maternal odor modulates both behavioral and electrophysiological responses to facial information in young infants (Durand et al., 2013, 2020; Endevelt-Shapira et al., 2021; Jessen, 2020). The mother’s body odor is a salient chemosensory stimulus that accompanies infants from the very beginning of life and promotes specific physiological, behavioral and neural responses already in neonates (Schaal et al., 2020 for review). By stimulating eye opening at birth (Doucet et al., 2007), maternal odor favors optimal exposure to the mother’s face during nursing and caregiving. Later on, infants are often carried by their mother while engaged in social interactions, leading to co-exposure to the mother’s odor and to various (un)familiar faces. This repeated co-occurrence of both sensory stimulations in the infants’ environment could generate a relevant association, leading to intersensory facilitation, as previously observed for audiovisual associations (e.g. Hyde et al., 2011; Lewkowicz et al., 2010; Neil et al., 2006).

Interestingly, compared to auditory and visual perception, odor perception is less sensitive to spatiotemporal precision (Sela & Sobel, 2010). This reduced spatiotemporal constraint could make odor cues particularly prone to foster the development of visual categorization by supporting the generalization of variable inputs into a single category. At the neural level, a dedicated connectivity could be shaped by the recurrent association between the mother’s odor and faces, so that the former would be able to boost the sensitivity of the face-selective network through reentrant signaling (Edelman, 1993). Such connectivity is supported by a body of research in adults revealing that the sole presentation of odors activates the right occipital cortex (e.g., Royet et al., 2001), and body odors in particular elicit neural activity in the lateral fusiform gyrus (e.g., W. Zhou & Chen, 2008), which is part of the face-selective network and is also functionally connected to the primary olfactory cortex (G. Zhou et al., 2019). Finally, it is also important to note that the odor effect is not driven by a mere enhancement of arousal or attention, as the general response to the rapid stream of visual stimulation remains unaffected by the presence of the mother’s body odor (Leleu et al., 2020; Rekow et al., 2020, 2021). This dissociation between the face-selective and general visual responses as a function of age reminds that the frequency-tagging approach allows to isolate and characterize distinct responses at different frequencies simultaneously within the same stimulation sequence.

### Olfactory-to-visual facilitation declines gradually as a function of age

Importantly, according to our main hypothesis, we found a gradual decline of the maternal odor effect as a function of age. The odor effect observed at the right occipital electrode O2 is the strongest for the youngest infants, and progressively decreases and reverses as age increases, such that the face-selective response becomes restricted to occipito-temporal sites in the presence of the mother’s body odor for the oldest infants. This supports prior studies in the audiovisual domain reporting intersensory facilitation in younger but not in older infants when unisensory perception has improved (Bahrick et al., 2004; Bahrick & Lickliter, 2000, 2004), and extends them to olfactory-visual interactions. Such developmental trade-off between olfaction and vision for efficient categorization relates to the inverse effectiveness principle whereby multisensory integration decreases as unisensory responses increases (e.g., Meredith & Stein, 1983; Regenbogen et al., 2016; Stevenson et al., 2012). This presumably stems from the fact that a key function of multisensory integration is the disambiguation of otherwise ambiguous unisensory events (Ernst & Bülthoff, 2004), as already reported in infants (Phillips-Silver & Trainor, 2005; Scheier et al., 2003), or in adults for odor effects on the perception of ambiguous facial expressions (e.g., Forscher & Li, 2012; Leleu et al., 2015; Poncet et al., 2021; W. Zhou & Chen, 2009). Similarly, in adults, a composite body odor pooled from 8 unfamiliar donors favors the categorization of ambiguous face-like objects while it does not influence human face categorization (Rekow, Baudouin, Durand, et al., 2022), which is readily achieved without another sensory cue and under tight visual constraints in the adult brain (e.g., Retter et al., 2020). Hence, the presence of maternal odor cues may help the infant brain to disambiguate inputs to reach optimal face categorization, unless visual inputs are sufficient to elicit a robust percept as face perceptual abilities improve over the first year of life. Future studies should investigate this interpretation by increasing, for instance, face categorization demand at 12 months of age, to determine whether intersensory facilitation from the mother’s odor reappears, as previously observed with audiovisual stimulations (Bahrick et al., 2010).

Alternatively, during development, maternal odor may become less associated with the face category in general. Early on, infants are already able to differentiate their own from another mother’s odor (e.g., Cernoch & Porter, 1985; Russell, 1976; Schaal et al., 1998). Thus, as face perceptual abilities develop, the mother’s body odor may be specifically associated with the recognition of her face, while the association with other faces weakens. In the same vein, the femininity of the mother’s odor may gradually become recognized by older infants and selectively associated with the categorization of female faces. This putative developmental shift is in line with previous evidence in the audiovisual domain that multisensory perception is broadly tuned during the first months and then gradually narrows toward more specific associations (Murray et al., 2016 for review). Maternal odor could even be progressively dissociated from the perception of faces, as the proportion of faces in the infant’s visual environment linearly decreases during the first year while the proportion of other body parts, especially hands, increases (Fausey et al., 2016). However, accumulating evidence shows that human odors are still associated with faces in adulthood (see Damon et al., 2021 for a recent review), indicating that body odor influence on face perception is not restricted to the maternal odor effect in the youngest infants.

Another non-mutually exclusive interpretation could be that maternal odor progressively loses the ability to trigger face categorization because its composition changes between 4 and 12 months. The mother’s body odor is a mixture of several compounds conveying a variety of cues about her traits (e.g., identity, femininity, humanity) and states (e.g., maternity, emotions). In particular, certain cues emanating from milk can be discriminated by infants, who differentiate human from non-human milk (Marlier & Schaal, 2005), lactating from non-lactating women (Makin & Porter, 1989), or early from late lactation milk (Klaey-Tassone et al., 2020). Given that young infants are generally more often breastfed than older infants, feeding status is a candidate factor for the maternal odor effect. However, when adding the feeding factor to the analysis, there is no evidence for an interaction between feeding and age, and no interaction with the odor effect (Table S6 and Figure S3, also for the absence of significant interactions with the infants’ sex). For other cues, such as the familiarity of the odor, there is no clear evidence in the literature, as previous infant studies about odor-face interactions that compared the own and another mother’s odor did not systematically evidence a comparable effect (Durand et al., 2020; Jessen, 2020). Finally, as mentioned above, a composite body odor pooled from males and nulliparous females is able to foster face perception in the adult brain (Rekow, Baudouin, Durand, et al., 2022; see also Cecchetto et al., 2020; Wudarczyk et al., 2016 for other non-maternal body odors), revealing the potency of a human odor beyond maternal cues, at least in adulthood. Therefore, future studies should carefully characterize the chemical profile of the mother’s odor as a function of age, and delineate which cues carried by this odor facilitate face categorization in the youngest infants.

### Conclusions

To conclude, we have shown that the mother’s odor effect on neural face categorization previously reported at 4 months (Leleu et al., 2020) gradually decreases with age as face-selective activity amplifies. This suggests that face categorization relies on maternal odor cues in developing infants until the visual system becomes able to readily achieve categorization by itself, generalizing previous evidence in the audiovisual domain that intersensory facilitation declines as unisensory perception improves (Bahrick et al., 2002; Bahrick & Lickliter, 2000, 2004). From a broad developmental perspective, our findings favor a multisensory view of knowledge acquisition (Gibson, 1969). However, this does not mean that perception strictly develops from multisensory to unisensory abilities, a large body of research having shown intersensory facilitation in adults when unisensory inputs are scarce or ambiguous (e.g., for odor-face association Forscher & Li, 2012; Leleu et al., 2015; Poncet et al., 2021; Rekow, Baudouin, Durand, et al., 2022). We thus support a lifespan view of multisensory perception (Bahrick & Lickliter, 2012; Murray et al., 2016; Lewkowicz & Bremner, 2020 for reviews), in which intersensory facilitation at any age is a function of how unisensory perception is effective on its own. In this context, early infancy would represent a key period of multisensory development, when the perceptual system is still immature and highly naïve toward the environment, while progressive maturation and experience would then reduce the need to bind multisensory cues for effective categorization in the environment.

## Acknowledgment

This work was financially supported by the French “Investissements d’Avenir” program, project ISITE-BFC (contract ANR-15-IDEX-0003), the “Conseil Régional Bourgogne Franche-Comté”, the FEDER (European Funding for Regional Economic Development), the French National Research Agency (contract ANR-19-CE28-0009), and the European Research Council (contract HUMANFACE 101055175). The authors are grateful to the participating parents and infants. They also thank Sylviane Martin for her help in recruiting them and Vincent Gigot for his assistance with EEG preprocessing.

## Notes

**Conflict of interest:** None.

### Competing Interest Statement

The authors have declared no competing interest.

