## Supplementary material for "Olfactory-to-visual facilitation in the infant brain declines gradually from 4 to 12 months"

### Supplementary data

**Supplementary method: EEG preprocessing (references are appended at the**

EEG data were filtered using a Butterworth filter (highpass filter, cutoff: 0.1 Hz, 4th order) and resampled to 200 Hz. Next, sequences were cropped from the beginning of the fade-in into 36-s segments (i.e., adding 2 s after the end of the sequence). The *Artifact Blocking* algorithm (Fujioka et al., 2011; Mourad et al., 2007) was applied to individual epochs with a correction threshold of ± 250 µV. Epochs were then segmented again from the end of the fade-in (i.e., from 1.833 s, corresponding to the first image of the full-contrast segment) until the end of the fade-out, i.e., in 32-s-long segments, and datasets were re-referenced according to a common average reference.

Following previous frequency-tagging EEG studies in infants (e.g., Leleu et al., 2020; Rekow et al., 2021, 2020), data were screened using two data-driven criteria to remove unusable epochs at an individual level and increase signal-to-noise ratio. The first criterion consisted in identifying epochs with no general visual response at 6 Hz and its second harmonic (12 Hz), used as a marker of the infant's attention to the stimulation. Baseline-corrected amplitude spectra were extracted for individual epochs and *Z*-scores were calculated (see Frequency-domain analysis) for medial occipital electrodes (POz, Oz, O1, O2) that typically exhibit the largest general visual response to a rapid stream of natural images (Leleu et al., 2020). Epochs failing to present a significant ($Z > 1.64$, $p < .05$, one-tailed, signal > noise) general visual response for 2 electrodes, at least, were removed. The second criterion consisted in identifying atypical epochs according to the scalp-wide power of the response at 1 Hz. For each epoch, the root mean square (RMS) amplitude across electrodes was calculated. Epochs were rejected if their RMS amplitude was ±2 SDs of the mean of all epochs.

**Table S1. Harmonic significance for the general visual response and the face-selective response.** For both responses, harmonic significance was estimated using *Z*-scores calculated on the average of all electrodes, participants and odor conditions. Harmonics were considered significant if their *Z*-score was > 1.64 ($p < .05$, one-tailed, signal > noise). For the face-selective response, the 6th harmonic was not considered as it corresponds to the 1st harmonic of the general visual response. Asterisks indicate significance (* $p < .05$, *** $p < .001$).

| | General visual response | | Face-selective response | |
|---|---|---|---|---|
| **Harmonic** | **Frequency (Hz)** | ***Z*-score** | **Frequency (Hz)** | ***Z*-score** |
| 1 | 6 | 148*** | 1 | 6.62*** |
| 2 | 12 | 167*** | 2 | 5.35*** |
| 3 | 18 | 80.8*** | 3 | 3.66*** |
| 4 | 24 | 49.8*** | 4 | 3.67*** |
| 5 | 30 | 8.97*** | 5 | 6.22*** |
| 6 | 36 | 5.07*** | | |
| 7 | 42 | 0.56 | 7 | 2.15* |
| 8 | 48 | -0.06 | 8 | -1.04 |

**Table S2. Electrode significance for the general visual response and the face-selective response.** For both responses, electrode significance was estimated using *Z*-scores calculated on responses summed across significant harmonics and averaged across participants and odor conditions. Electrodes were considered significant if their *Z*-score was > 2.93 (*p* < .05, one-tailed, signal > noise, Bonferroni-corrected for 30 electrodes). *Z*-scores are presented in decreasing order and asterisks indicate significance (* *p* < .05, ** *p* < .01, *** *p* < .001). Electrodes highlighted in grey were used for further analyses.

| | General visual response | | Face-selective response | |
|---|---|---|---|---|
| **Rank** | **Electrode** | ***Z*-score** | **Electrode** | ***Z*-score** |
| 1 | **Oz** | **168*** ** | **P8** | **19.5*** ** |
| 2 | **O2** | **133*** ** | **P7** | **13.1*** ** |
| 3 | **POz** | **122*** ** | Oz | 6.16*** |
| 4 | **O1** | **102*** ** | POz | 5.32*** |
| 5 | P3 | 71.1*** | **O1** | **5.31*** ** |
| 6 | CP6 | 61.0*** | **O2** | **4.38*** ** |
| 7 | FC2 | 46.9*** | CP2 | 4.25*** |
| 8 | CP5 | 45.1*** | P3 | 3.45** |
| 9 | P4 | 43.1*** | P4 | 3.38* |
| 10 | P7 | 39.6*** | F7 | 3.27* |
| 11 | CP2 | 33.4*** | T7 | 3.21* |
| 12 | FC1 | 32.4*** | FC6 | 2.70 |
| 13 | Fz | 31.6*** | FC1 | 2.63 |
| 14 | T8 | 29.6*** | CP5 | 2.60 |
| 15 | CP1 | 28.9*** | FC2 | 2.55 |
| 16 | Pz | 28.9*** | Pz | 2.51 |
| 17 | FC5 | 28.7*** | CP1 | 2.39 |
| 18 | F3 | 27.7*** | F8 | 2.07 |
| 19 | F4 | 26.9*** | F4 | 2.05 |
| 20 | P8 | 26.8*** | Cz | 1.92 |
| 21 | C3 | 26.7*** | T8 | 1.81 |
| 22 | Fp2 | 26.5*** | CP6 | 1.77 |
| 23 | F7 | 26.0*** | F3 | 1.76 |
| 24 | FC6 | 25.9*** | FC5 | 1.71 |
| 25 | Cz | 25.1*** | C3 | 1.61 |
| 26 | C4 | 24.3*** | Fz | 1.19 |
| 27 | F8 | 21.4*** | C4 | 1.07 |
| 28 | Fpz | 20.5*** | Fpz | 0.65 |
| 29 | T7 | 20.3*** | Fp2 | 0.53 |
| 30 | Fp1 | 17.7*** | Fp1 | -0.13 |

**Table S3. ANCOVA on the amplitude of the face-selective response in the baseline odor context.** A first analysis was conducted in the baseline odor context to determine the effect of AGE and its interactions with the other factors. Significant effects are reported in red. Greenhouse-Geisser epsilon ($\varepsilon$) and corresponding adjusted *p*-values are reported when sphericity was violated (as estimated by Mauchly's test, unreported). The significant HARM × AGE interaction is decomposed in the effect of AGE for each harmonic separately. Another ANCOVA was run on the response summed across the two first harmonics (Harmonics 1 + 2). HARM: Harmonic, ELEC: Electrode, SS: sum of square, Df: degree of freedom, MS: mean square, GG: Greenhouse-Geisser, Adj: adjusted.

| | SS | Df | MS | *F* | *p* | $\eta_p^2$ | GG $\varepsilon$ | GG Adj *p* |
|---|---|---|---|---|---|---|---|---|
| AGE | 21.79 | 1 | 21.79 | 8.41 | .006 | .149 | | |
| Error | 124.43 | 48 | 2.59 | | | | | |
| HARM*AGE | 39.18 | 5 | 7.84 | 5.54 | .000 | .104 | .38 | .006 |
| Error | 339.28 | 240 | 1.41 | | | | | |
| ELEC*AGE | 6.42 | 3 | 2.14 | 2.08 | .106 | .041 | .80 | .121 |
| Error | 148.31 | 144 | 1.03 | | | | | |
| HARM*ELEC*AGE | 10.17 | 15 | 0.68 | 0.79 | .686 | .016 | .25 | .523 |
| Error | 614.90 | 720 | 0.85 | | | | | |
| ***Harmonic 1*** | | | | | | | | |
| AGE | 38.85 | 1 | 38.85 | 6.03 | .018 | .112 | | |
| Error | 309.33 | 48 | 6.44 | | | | | |
| ***Harmonic 2*** | | | | | | | | |
| AGE | 19.93 | 1 | 19.93 | 12.01 | .001 | .200 | | |
| Error | 79.63 | 48 | 1.66 | | | | | |
| ***Harmonic 3*** | | | | | | | | |
| AGE | 0.62 | 1 | 0.62 | 1.83 | .182 | .037 | | |
| Error | 16.12 | 48 | 0.34 | | | | | |
| ***Harmonic 4*** | | | | | | | | |
| AGE | 0.09 | 1 | 0.09 | 0.22 | .638 | .005 | | |
| Error | 19.94 | 48 | 0.42 | | | | | |
| ***Harmonic 5*** | | | | | | | | |
| AGE | 1.49 | 1 | 1.49 | 2.17 | .148 | .043 | | |
| Error | 32.89 | 48 | 0.69 | | | | | |
| ***Harmonic 7*** | | | | | | | | |
| AGE | 0.00 | 1 | 0.00 | 0.00 | .996 | .000 | | |
| Error | 5.80 | 48 | 0.12 | | | | | |
| ***Harmonics 1 + 2*** | | | | | | | | |
| AGE | 114.42 | 1 | 114.42 | 11.17 | .002 | .189 | | |
| Error | 491.80 | 48 | 10.25 | | | | | |
| ELEC*AGE | 20.36 | 3 | 6.79 | 1.51 | .215 | .030 | .78 | .224 |
| Error | 648.71 | 144 | 4.50 | | | | | |

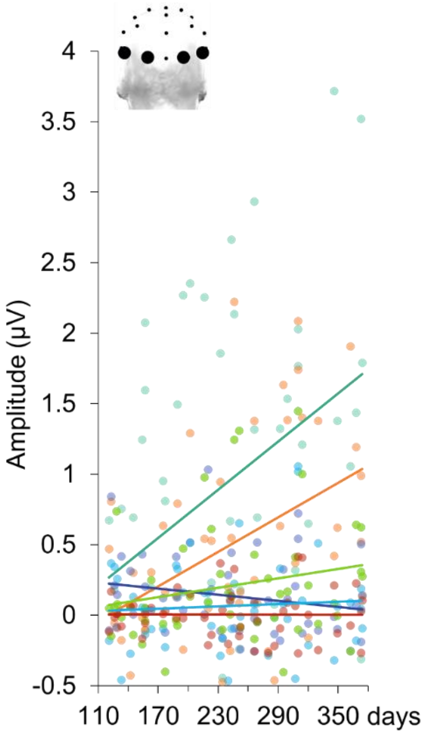


**Figure S1.** Individual amplitudes of the face-selective response recorded in the baseline odor context over the occipital temporal electrodes for each harmonic separately (see Tables S3 for color codes) as a function of age. Each dot represents an infant and each line represents the linear regression for a harmonic. The effect of *Age* is significant for the two first harmonics (dark green and orange).

**Table S4. ANCOVA on the amplitude of the face-selective response in both odor contexts.** A second analysis was conducted to determine the effect of ODOR, its interaction with AGE and their interactions with the other factors. Significant effects are reported in red (one effect is reported in orange because significance did not survive the Greenhouse-Geisser correction). Greenhouse-Geisser epsilon ($\varepsilon$) and corresponding adjusted $p$-values are reported when sphericity was violated (as estimated by Mauchly's test, unreported). The significant HARM × ODOR × ELEC × AGE interaction is decomposed in the effects involving these factors for each harmonic separately. HARM: Harmonic, ELEC: Electrode, SS: sum of square, Df: degree of freedom, MS: mean square, GG: Greenhouse-Geisser, Adj: adjusted.

| | SS | Df | MS | $F$ | $p$ | $\eta_p^2$ | GG $\varepsilon$ | GG Adj $p$ |
|---|---|---|---|---|---|---|---|---|
| ODOR | 1.40 | 1 | 1.40 | 1.48 | .233 | .029 | | |
| ODOR*AGE | 0.77 | 1 | 0.77 | 0.80 | .376 | .016 | | |
| Error | 46.01 | 48 | 0.96 | | | | | |
| HARM*ODOR | 12.49 | 5 | 2.50 | 2.22 | .053 | .044 | .46 | .105 |
| HARM*ODOR*AGE | 14.89 | 5 | 2.98 | 2.65 | .025 | .052 | .46 | .067 |
| Error | 269.43 | 240 | 1.12 | | | | | |
| ODOR*ELEC | 2.65 | 3 | 0.88 | 1.26 | .290 | .026 | | |
| ODOR*ELEC*AGE | 2.05 | 3 | 0.68 | 0.98 | .406 | .020 | | |
| Error | 100.88 | 144 | 0.70 | | | | | |
| HARM*ODOR*ELEC | 33.76 | 15 | 2.25 | 3.58 | .000 | .069 | .34 | .004 |
| HARM*ODOR*ELEC*AGE | 40.38 | 15 | 2.69 | 4.28 | .000 | .082 | .34 | .001 |
| Error | 452.41 | 720 | 0.63 | | | | | |
| ***Harmonic 1*** | | | | | | | | |
| ODOR | 7.74 | 1 | 7.74 | 2.05 | .159 | .041 | | |
| ODOR*AGE | 4.90 | 1 | 4.90 | 1.30 | .260 | .026 | | |
| Error | 181.27 | 48 | 3.78 | | | | | |
| ODOR*ELEC | 32.48 | 3 | 10.83 | 4.32 | .006 | .083 | | |
| ODOR*ELEC*AGE | 37.68 | 3 | 12.56 | 5.01 | .002 | .095 | | |
| Error | 361.01 | 144 | 2.51 | | | | | |
| ***Harmonic 2*** | | | | | | | | |
| ODOR | 4.77 | 1 | 4.77 | 3.52 | .067 | .068 | | |
| ODOR*AGE | 8.30 | 1 | 8.30 | 6.12 | .017 | .113 | | |
| Error | 65.05 | 48 | 1.36 | | | | | |
| ODOR*ELEC | 1.02 | 3 | 0.34 | 0.57 | .637 | .012 | .76 | .592 |
| ODOR*ELEC*AGE | 1.35 | 3 | 0.45 | 0.75 | .522 | .015 | .76 | .490 |
| Error | 86.25 | 144 | 0.60 | | | | | |
| ***Harmonic 3*** | | | | | | | | |
| ODOR | 0.25 | 1 | 0.25 | 0.52 | .474 | .011 | | |
| ODOR*AGE | 0.67 | 1 | 0.67 | 1.37 | .248 | .028 | | |
| Error | 22.34 | 48 | 0.49 | | | | | |
| ODOR*ELEC | 1.61 | 3 | 0.54 | 2.14 | .097 | .043 | .84 | .108 |
| ODOR*ELEC*AGE | 1.81 | 3 | 0.60 | 2.40 | .070 | .048 | .84 | .082 |
| Error | 36.14 | 144 | 0.25 | | | | | |
| ***Harmonic 4*** | | | | | | | | |
| ODOR | 0.11 | 1 | 0.11 | 0.32 | .572 | .007 | | |
| ODOR*AGE | 0.33 | 1 | 0.67 | 1.01 | .319 | .021 | | |
| Error | 15.69 | 48 | 0.33 | | | | | |
| ODOR*ELEC | 0.89 | 3 | 0.30 | 1.06 | .369 | .022 | | |
| ODOR*ELEC*AGE | 1.19 | 3 | 0.40 | 1.42 | .239 | .029 | | |
| Error | 40.26 | 144 | 0.28 | | | | | |

| ***Harmonic 5*** | SS | Df | MS | *F* | *p* | $\eta_p^2$ | GG $\varepsilon$ | GG Adj *p* |
|---|---|---|---|---|---|---|---|---|
| ODOR | 0.95 | 1 | 0.95 | 1.85 | .180 | .037 | | |
| ODOR*AGE | 1.28 | 1 | 1.28 | 2.49 | .121 | .049 | | |
| Error | 24.72 | 48 | 0.52 | | | | | |
| ODOR*ELEC | 0.18 | 3 | 0.06 | 0.38 | .770 | .008 | .83 | .732 |
| ODOR*ELEC*AGE | 0.11 | 3 | 0.04 | 0.23 | .873 | .005 | .83 | .838 |
| Error | 22.47 | 144 | 0.16 | | | | | |
| ***Harmonic 7*** | | | | | | | | |
| ODOR | 0.07 | 1 | 0.07 | 0.62 | .434 | .013 | | |
| ODOR*AGE | 0.18 | 1 | 0.18 | 1.60 | .211 | .032 | | |
| Error | 5.37 | 48 | 0.11 | | | | | |
| ODOR*ELEC | 0.23 | 3 | 0.08 | 1.53 | .208 | .031 | | |
| ODOR*ELEC*AGE | 0.29 | 3 | 0.10 | 1.92 | .129 | .038 | | |
| Error | 7.17 | 144 | 0.05 | | | | | |
| ***Harmonics 1 + 2*** | | | | | | | | |
| ODOR | 24.65 | 1 | 24.65 | 5.15 | .028 | .097 | | |
| ODOR*AGE | 25.95 | 1 | 25.95 | 5.43 | .024 | .102 | | |
| Error | 229.57 | 48 | 4.78 | | | | | |
| ODOR*ELEC | 27.58 | 3 | 9.19 | 3.15 | .027 | .061 | | |
| ODOR*ELEC*AGE | 29.40 | 3 | 9.80 | 3.35 | .021 | .065 | | |
| Error | 420.95 | 144 | 2.92 | | | | | |
| ***Electrode O2*** | | | | | | | | |
| ODOR | 48.43 | 1 | 48.43 | 21.56 | .000 | .310 | | |
| ODOR*AGE | 49.05 | 1 | 49.05 | 21.83 | .000 | .313 | | |
| Error | 107.84 | 48 | 2.25 | | | | | |
| ***Electrode P8*** | | | | | | | | |
| ODOR | 1.27 | 1 | 1.27 | 0.29 | .596 | .006 | | |
| ODOR*AGE | 0.00 | 1 | 0.00 | 0.00 | .999 | .000 | | |
| Error | 213.12 | 48 | 4.44 | | | | | |
| ***Electrode O1*** | | | | | | | | |
| ODOR | 2.46 | 1 | 2.46 | 0.68 | .413 | .014 | | |
| ODOR*AGE | 5.65 | 1 | 5.65 | 1.57 | .216 | .032 | | |
| Error | 172.45 | 48 | 3.59 | | | | | |
| ***Electrode P7*** | | | | | | | | |
| ODOR | 0.08 | 1 | 0.08 | 0.02 | .879 | .000 | | |
| ODOR*AGE | 0.65 | 1 | 0.65 | 0.20 | .658 | .004 | | |
| Error | 157.11 | 48 | 3.27 | | | | | |

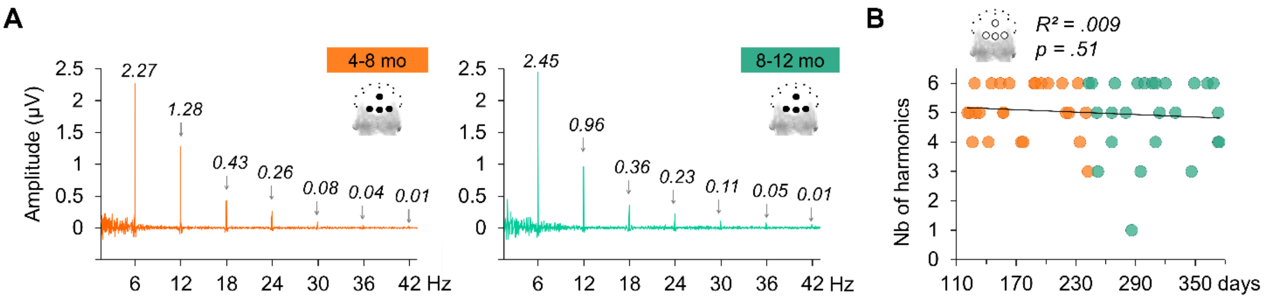


**Figure S2. The development of the general visual response with age. A.** Amplitude spectra recorded in the baseline odor context for the youngest (4-8 months, orange) and the oldest (8-12 months, green) infants averaged for the 4 middle occipital electrodes (POz, Oz, O1, O2). **B.** Maximum number of significant harmonics recorded among the 4 middle occipital electrodes as a function of age. Each color circle represents an individual infant data depending on its subgroup (orange: 4-8 months, green: 8-12 months).

**Table S5. ANCOVAs on the amplitude of the general visual response.** A first analysis was conducted in the baseline odor context to determine the effect of AGE and its interactions with the other factors. A second analysis was conducted to determine the effect of ODOR, its interaction with AGE and their interactions with the other factors. This analysis was also conducted after removing one infant who had an outlying odor effect. Greenhouse-Geisser epsilon ($\varepsilon$) and corresponding adjusted *p*-values are reported when sphericity was violated (as estimated by Mauchly's test, unreported). HARM: Harmonic, ELEC: Electrode, SS: sum of square, Df: degree of freedom, MS: mean sum, GG: Greenhouse Geiser, Adj: adjusted.

| ***Baseline odor context*** | SS | Df | MS | *F* | *p* | $\eta_p^2$ | *GG ε* | GG Adj *p* |
|---|---|---|---|---|---|---|---|---|
| AGE | 1.97 | 1 | 1.97 | 0.32 | .575 | .007 | | |
| Error | 297.19 | 48 | 6.19 | | | | | |
| HARM*AGE | 14.62 | 5 | 2.92 | 0.94 | .454 | .019 | .23 | .348 |
| Error | 743.98 | 240 | 3.10 | | | | | |
| ELEC*AGE | 4.71 | 3 | 1.57 | 1.89 | .134 | .038 | .61 | .160 |
| Error | 119.50 | 144 | 0.83 | | | | | |
| HARM*ELEC*AGE | 5.32 | 15 | 0.35 | 1.03 | .419 | .021 | .14 | .362 |
| Error | 247.35 | 720 | 0.34 | | | | | |
| ***Both odor contexts*** | | | | | | | | |
| ODOR | 0.40 | 1 | 0.40 | 0.27 | .606 | .006 | | |
| ODOR*AGE | 0.11 | 1 | 0.11 | 0.07 | .791 | .001 | | |
| Error | 72.10 | 48 | 1.50 | | | | | |
| HARM*ODOR | 0.33 | 5 | 0.07 | 0.11 | .991 | .002 | .23 | .786 |
| HARM*ODOR*AGE | 0.09 | 5 | 0.02 | 0.03 | .999 | .001 | .23 | .897 |
| Error | 147.54 | 240 | 0.61 | | | | | |
| ODOR*ELEC | 0.13 | 3 | 0.04 | 0.28 | .842 | .006 | .78 | .792 |
| ODOR*ELEC*AGE | 0.16 | 3 | 0.05 | 0.35 | .790 | .007 | .78 | .739 |
| Error | 21.59 | 144 | 0.15 | | | | | |
| HARM*ODOR*ELEC | 0.16 | 15 | 0.01 | 0.14 | .999 | .003 | .19 | .928 |
| HARM*ODOR*ELEC*AGE | 0.51 | 15 | 0.03 | 0.45 | .962 | .009 | .19 | .708 |
| Error | 54.31 | 720 | 0.08 | | | | | |
| ***Both odor contexts without 1 outlier*** | | | | | | | | |
| ODOR | 0.62 | 1 | 0.62 | 0.97 | .329 | .020 | | |
| ODOR*AGE | 0.61 | 1 | 0.61 | 0.96 | .333 | .020 | | |
| Error | 30.13 | 47 | 0.64 | | | | | |
| HARM*ODOR | 0.59 | 5 | 0.12 | 0.40 | .848 | .008 | .27 | .591 |
| HARM*ODOR*AGE | 0.53 | 5 | 0.11 | 0.36 | .875 | .008 | .27 | .615 |
| Error | 68.70 | 235 | 0.29 | | | | | |
| ODOR*ELEC | 0.13 | 3 | 0.04 | 0.30 | .828 | .006 | .74 | .768 |
| ODOR*ELEC*AGE | 0.13 | 3 | 0.04 | 0.30 | .827 | .006 | .74 | .767 |
| Error | 20.23 | 141 | 0.14 | | | | | |
| HARM*ODOR*ELEC | 0.16 | 15 | 0.01 | 0.15 | .999 | .003 | .18 | .918 |
| HARM*ODOR*ELEC*AGE | 0.46 | 15 | 0.03 | 0.42 | .974 | .008 | .18 | .722 |
| Error | 51.86 | 705 | 0.07 | | | | | |

**Table S6. ANCOVA on the amplitude of the face-selective response at electrode O2 including the SEX and FEEDING factors.** An analysis was conducted to determine whether the ODOR effect and the interaction ODOR × AGE observed at O2 in the main analysis remain when the between-subject categorical factors SEX (26 females *vs.* 24 males) and FEEDING (17 breastfed *vs*. 33 bottle-fed infants) are added. Both effects remained significant (reported in red). In addition, the AGE × SEX, AGE × FEEDING, ODOR × AGE × SEX, and ODOR × AGE × FEEDING interactions were not significant, indicating that sex and feeding status were not significantly different as a function of age (young infants were not significantly more (fe)males or more breastfed than older infants) and did not contribute to the maternal odor effect observed for the youngest infants. SS: sum of square, Df: degree of freedom, MS: mean square.

| ***Electrode O2*** | SS | Df | MS | *F* | *p* | $\eta_p^2$ |
|---|---|---|---|---|---|---|
| ODOR | 49.33 | 1 | 49.33 | 21.26 | .000 | .316 |
| ODOR*AGE | 48.74 | 1 | 48.74 | 21.01 | .000 | .314 |
| ODOR*AGE*SEX | 0.66 | 1 | 0.66 | 0.28 | .597 | .006 |
| ODOR*AGE*FEEDING | 0.23 | 1 | 0.23 | 0.10 | .753 | .002 |
| Error | 106.73 | 46 | 2.32 | | | |
| AGE*SEX | 0.05 | 1 | 0.05 | 0.01 | .914 | .000 |
| AGE*FEEDING | 0.47 | 1 | 0.47 | 0.11 | .745 | .002 |
| Error | 200.90 | 46 | 4.37 | | | |

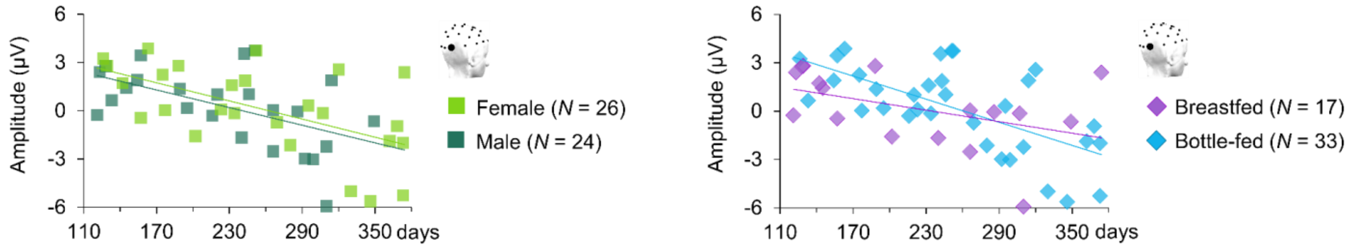


**Figure S3. Maternal odor effect at electrode O2 as a function of age and sex (left) or feeding status (right).** Each square or diamond represents the odor effect (amplitude of the face-selective response in the maternal minus the baseline odor context) for an individual infant depending on their sex (light green: female, *N* = 26; dark green: male, *N* = 24) or feeding status (purple: breastfed, *N* = 17; blue: bottle-fed, *N* = 33).
